# Genetic landscape of Human neutrophil antigen variants in India from population-scale genomes

**DOI:** 10.1101/2022.06.23.497282

**Authors:** Mercy Rophina, Rahul C Bhoyar, Mohamed Imran, Vigneshwar Senthivel, Mohit Kumar Divakar, Anushree Mishra, Bani Jolly, Sridhar Sivasubbu, Vinod Scaria

## Abstract

**Background:** Antibodies against human neutrophil antigens (HNAs) play a significant role in various clinical conditions such as neonatal alloimmune neutropenia (NAIN). Transfusion-related acute lung injury (TRALI) and other non-hemolytic transfusion reactions. This study aims to identify the genotype and allele frequencies of HNAs in the healthy Indian population.

**Methods:** Genetic variants from whole genomes of 1029 healthy Indian individuals were retrieved to accurately perform frequency estimation of HNA-1, HNA-3, HNA-4 and HNA-5 alleles using in-house computational pipeline.

**Results:** In HNA class I, the genotype frequencies of FCGR3B*01 (HNA1a/a), FCGR3B*02 (HNA1b/b) and FCGR3B*03 (HNA1c/c) were 0.29%, 27.31% and 1.75% respectively. In HNA-3 the frequencies of HNA3a/a (SLC44A2*01), HNA3a/b and HNA3b/b (SLC44A2*02) were found to be 62.0%, 31.7% and 5.8% respectively. Frequency of ITGAM*01 encoding HNA4a/a was 90.1% and that of ITGAM*02 encoding HNA4b/b was 0.3%. Furthermore, HNA5a/a (ITGAL*01) and HNA5b/b (ITGAL*02) were found to have 12.9% and 48.6% genotype frequencies in the Indian population respectively. It was also found that the allele frequency HNA-5 variant, rs2230433 (ITGAL_chr16:30506720G>C) encoding 5b allele was highly prevalent (78.2%) in the Indian population which was comparable to South Asians (65.6%) but differed greatly from East Asians (14.3%), Latino Americans (25.7%), African-Americans (42.2%), European-Finnish (25.4%), European-non-Finnish (29.4%), Greater Middle Easterners (34.5%), Amish (30.2%) and Ashkenazi Jewish (31.4%).

**Conclusion:** This study presents the first comprehensive report of HNA variant and genotype frequencies using large scale representative whole genome sequencing data of the Indian population. Significant difference was observed in the prevalence of HNA5a and HNA5b in India in comparison to other global populations.

## Introduction

Human neutrophil antigens (HNAs), are a group of glycoprotein antigens that are expressed on the surface of human neutrophils^1,2^. These antigens are systematically classified into five major classes from HNA-1 to HNA-5 on the basis of serologically defined epitopes. Molecular characterization revealed that antigens of HNA class 1-5 are encoded by FCGR3B, CD177, CLT2, ITGAM and ITGAL genes respectively^2^. Although HNA-1 and HNA-2 have been found specifically expressed on the neutrophil surfaces,^3^ classes HNA-3 to HNA-5 have a broad range of expression in other blood cells^4^. HNAs play a significant clinical role in the field of transfusion and transplantation medicine as antibodies elicited against these antigens (both allo- and auto-) are implicated in the range of disease conditions such as transfusion-related acute lung injury (TRALI), neonatal alloimmune neutropenia (NAIN)^5,6,7,8^ autoimmune neutropenia (AIN)^9,10,11^ and febrile nonhemolytic transfusion reactions (FNHTR)^12^.

Owing to their clinical relevance, variant and genotype frequencies of HNAs have been studied among various global populations including Iranian^13^, Japanese^14^, Malays, Chinese^15^, Hong Kong^16^, Thai^17^, Turkish, German^18^, African American^19,20^, Danish and Zambian^21^.

While there is a paucity of reports on the genotype frequencies of HNA in the Indian population encompassing a sixth of the world population, the recent availability of population-scale genomic datasets have motivated us to undertake a systematic analysis of allele and genotype frequencies of HNA variants in Indian population. We hope our analysis would also provide insights and also potentially to estimate the risk of HNA alloimmunizations and devisee prevention strategies.

## Materials and Methods

### Reference and study datasets

A total of ten (10) variants in 4 human genes as approved by the International Society of Blood Transfusion - Granulocyte Immunobiology Working Party (ISBT-GIWP) for encoding alleles of human neutrophil antigens class I - V was used for the analysis. **Table 1** provides a comprehensive summary of these reference variants. The IndiGen dataset of genomic variants from whole genome sequences of 1029 healthy individuals from India was taken up for the analysis. Briefly, this dataset encompassed a total of 55,898,122 variants. The 10 variants associated with HNA were parsed in this dataset to retrieve the genotype and allele frequencies.. A schematic summary of the variant processing is shown in **Figure 1.**

**Table 1.** List of ISBT approved human neutrophil antigen variants used as reference in the study

| HNA Class | Chr | Pos (hg38) | RSID | Ref | Alt |
| --- | --- | --- | --- | --- | --- |
| HNA-1 | chr1 | 161629781 | rs2290834 | T | C |
| HNA-1 | chr1 | 161629853 | rs147574249 | T | C |
| HNA-1 | chr1 | 161629864 | rs5030738 | G | T |
| HNA-1 | chr1 | 161629903 | rs448740 | T | C |
| HNA-1 | chr1 | 161629983 | rs527909462 | A | G |
| HNA-1 | chr1 | 161629989 | rs200688856 | G | C |
| HNA-3 | chr19 | 10631490 | rs147820753 | C | T |
| HNA-3 | chr19 | 10631494 | rs2288904 | A | G |
| HNA-4 | chr16 | 31265490 | rs1143679 | G | A |
| HNA-5 | chr16 | 30506720 | rs2230433 | G | C |

**Figure 1.**
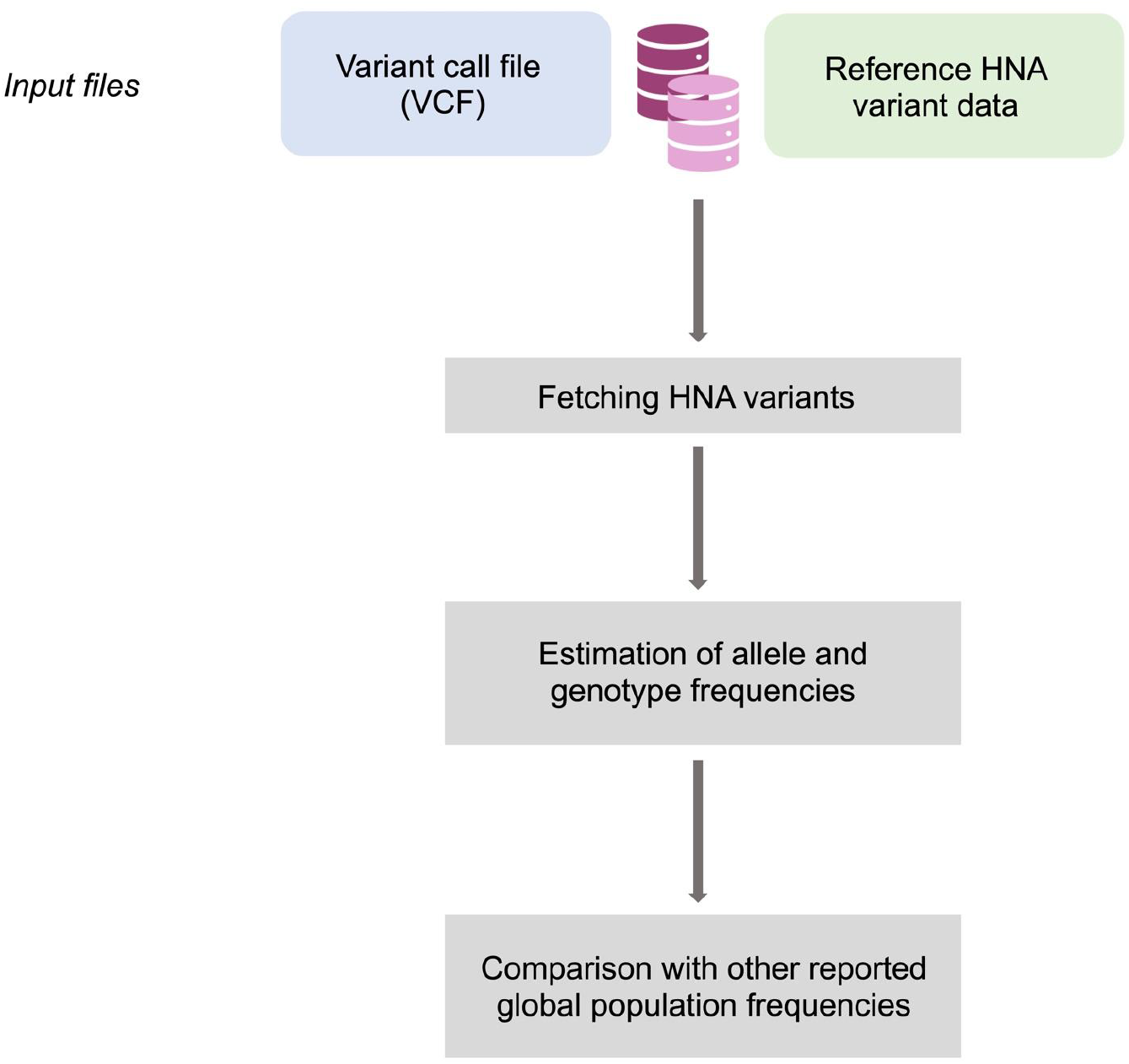
Schematic representation of the methodical workflow followed in the identification and estimation of HNA frequencies in India.

### Comparison of frequencies with other global populations

With the view of identifying significant differences in the prevalence of human neutrophil antigen variants in the Indian population, HNA allele frequencies were compared with major global population datasets retrieved from gnomAD database22. Fisher’s exact test with a p value <= 0.05 was used to estimate the statistical significance.

## Results

### Frequencies of HNA variants in Indian population

HNA variant frequencies were analyzed in a total of 1029 whole genome sequences. **Table 2** summarizes the allele frequencies of HNA variants in the Indian population. Based on the zygosity information the genotype frequencies were duly estimated for HNA-1, 3, 4 and 5 alleles. In HNA class I, genotype frequencies of HNA1a/a, HNA1b/b and HNA1c/c were observed as 0.29%, 27.31% and 1.75% respectively. Similarly, HNA3a/a, HNA3a/b and HNA3b/b of HNA class 3 were found at 62.00%, 31.68% and 5.83% respectively. In HNA class 4 and 5, the genotype frequencies were as follows, HNA4a/a - 90.18%, HNA4a/b - 9.23%, HNA4b/b - 0.29%, HNA5a/a - 12.93%, HNA5a/b - 38.10% and HNA5b/b - 48.59%.

**Table 2.** Frequencies of HNA variants observed in the Indian population

| HNA Class | Chr | Pos (hg38) | RSID | Ref | Alt | Allele count | Allele Number | Allele Frequency |
| --- | --- | --- | --- | --- | --- | --- | --- | --- |
| HNA-1 | chr1 | 161629781 | rs2290834 | T | C | AC=39 | AN=1070 | AF=0.036 |
| HNA-1 | chr1 | 161629853 | rs147574249 | T | C | AC=89 | AN=1186 | AF=0.075 |
| HNA-1 | chr1 | 161629864 | rs5030738 | G | T | AC=158 | AN=1210 | AF=0.131 |
| HNA-1 | chr1 | 161629903 | rs448740 | T | C | AC=1172 | AN=1402 | AF=0.836 |
| HNA-1 | chr1 | 161629983 | rs5279094<br>62 | A | G | AC=192 | AN=1662 | AF=0.116 |
| HNA-1 | chr1 | 161629989 | rs2006888<br>56 | G | C | AC=198 | AN=1670 | AF=0.119 |
| HNA-3 | chr19 | 10631490 | rs14782075<br>3 | C | T | NA | NA | NA |
| HNA-3 | chr19 | 10631494 | rs2288904 | A | G | AC=1602 | AN=2048 | AF=0.782 |
| HNA-4 | chr16 | 31265490 | rs1143679 | G | A | AC=101 | AN=2052 | AF=0.049 |
| HNA-5 | chr16 | 30506720 | rs2230433 | G | C | AC=1392 | AN=2050 | AF=0.679 |

### HNA variant frequencies among various global populations

Minor allele frequencies of HNA variants were fetched from the study dataset and were duly compared with global population frequencies. **Figure 2** provides a schematic overview of the distribution of the frequency distribution HNA variants in various populations. Comprehensive summary of genotype frequency comparison among globally reported populations is summarized in **Table 3.** Of the total, 5 variants were found to have significantly distinct differences (P <0.05) in Indian population in comparison to all the global populations. (FCGR3B_chr1:161629853T>C, FCGR3B_chr1:161629983A>G, FCGR3B_chr1:161629989G>C, ITGAM_chr16:31265490G>A and ITGAL_chr16:30506720G>C). The data is summarized in **Table 4.**

**Figure 2.**
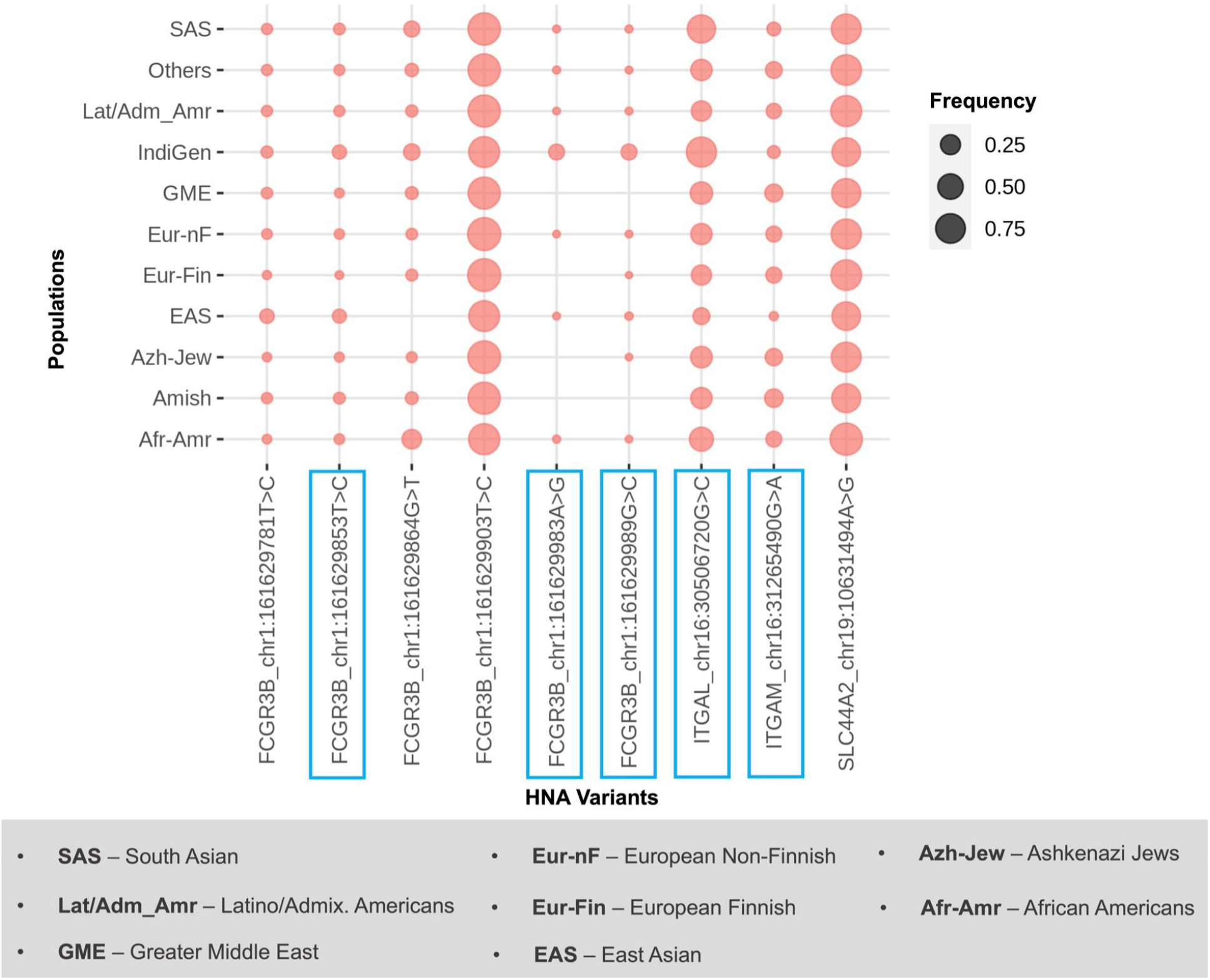
Schematic comparison of frequency distribution of ISBT approved human neutrophil antigen variants among global populations. Variants whose frequencies were found significantly distinct in India in comparison to all other global populations are marked in blue.

**Table 3.** HNA genotype frequencies observed among various global populations

| Genotype | India<br>(N=1029) | Hong<br>Kong<br>(N=300) | Guangzhou<br>(N=493) | British<br>(N=97) | German<br>(N=119) | Turkish<br>(N=118) | Danish<br>(N=200) | Zambians<br>(N=200) | Iranian<br>(N=150) | Japanese<br>(N=570) | Malays<br>(N=97) | Thai<br>(N=300) |
| --- | --- | --- | --- | --- | --- | --- | --- | --- | --- | --- | --- | --- |
| HNA-1<br>a/a | 3 | 137 | 210 | 11 | 0.151 | 0.161 | 26 | 39 | 14/150 | 193/570 | 0.186 | - |
| HNA-1<br>b/b | 281 | 28 | 44 | 60 | 0.361 | 0.305 | 84 | 32 | 43/150 | 64/570 | 0.134 | - |
| HNA-1<br>a/b |  | 133 | 238 | 65 | 0.479 | 0.517 | 78 | 49 | 37/150 | 266/570 | 0.68 | - |
| HNA-3<br>a/a | 60 | 154 | 104 | 79 | 0.555 | 0.559 | 245 | 183 | 38.7/150 | 238/570 | 0.041 | - |
| HNA-3<br>a/b | 326 | 118 | 80 | 57 | 0.378 | 0.356 | 106 | 10 | 48.6/150 | 269/570 | 0.918 | - |
| HNA-3<br>b/b | 638 | 28 | 11 | 4 | 0.067 | 0.085 | 15 | 0 | 12.7/150 | 63/570 | 0.041 | - |
| HNA-4<br>a/a | 928 | 297 | 489 | 110 | 0.832 | 0.763 | 163 | 146 | 73.3/150 | 570/570 | 1 | 284/300 |
| HNA-4<br>a/b | 95 | 3 | 4 | 27 | 0.151 | 0.237 | 44 | 31 | 24/150 | 0/570 | 0 | 16/300 |
| HNA-4<br>b/b | 3 | 0 | 0 | 3/97 | 0.017 | 0 | 3 | 4 | 2.7/150 | 0/570 | 0 | 0/300 |
| HNA-5<br>a/a | 133 | 215 | 361 | 76/97 | 0.555 | 0.585 | 111 | 42 | 51.3/150 | 364/570 | 0.443 | 186/300 |
| HNA-5<br>a/b | 392 | 81 | 120 | 54/97 | 0.353 | 0.339 | 82 | 105 | 40.7/150 | 125/570 | 0.495 | 86/300 |
| HNA-5<br>b/b | 500 | 4 | 12 | 10/97 | 0.092 | 0.076 | 17 | 42 | 8/150 | 19/570 | 0.062 | 18/300 |

**Table 4.**
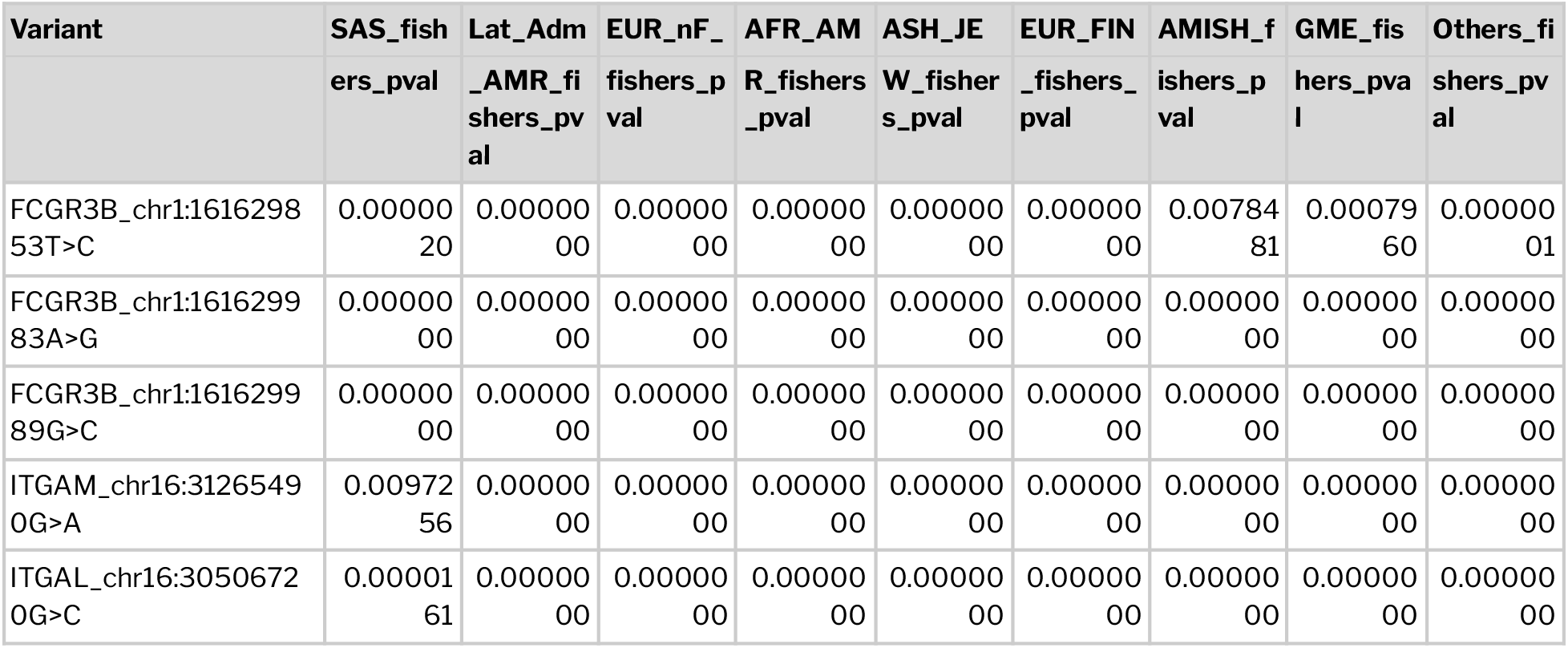
Summary of blood HNA variants found significantly distinct between India and global population datasets

## Discussion and Conclusion

Understanding the HNA variants are of clinical relevance due to their importance in allo and auto immunization resulting in diseases like TRALI, NAIN, AIN and graft rejections. It is therefore not surprising that the recent years have witnessed multiple attempts to decipher population specific HNA genotype and allele frequency profiles. While no population-scale datasets for HNA variants have been available for India, encompassing over a sixth of the world population, the recent availability of population-scale genome datasets motivated us to analyse allele and genotype frequencies of HNA-1a, -1b, -3a, -3b, -4a and -5a. Whole genome sequencing data of 1029 unrelated self declared healthy individuals was used to perform the *in-silico* genotyping analysis.

Alloimmunizations against HNA-1 systems has been well studied as a major cause of neonatal neutropenia and TRALI in Caucasian populations. Striking differences in prevalence of HNA-1 alleles have been observed in global populations with HNA-1a allele being the frequent in most Asian populations, whereas HNA-1b was found common in Caucasian^23^. A recent study by Hajar et al in 2019 suggested that unlike other Asian populations, HNA-1b was found common in samples with Indian ancestry^15^. In concordance with the earlier findings, HNA-1b/b homozygous individuals were found to constitute 27.3% of the population.

Allele encoding HNA-3a were found at a frequency of 0.78 in the Indian population which was concordant with the previous observations in Causian and Chinese populations with frequencies of 0.79 and 0.74 respectively ^24,20,23^. Antibodies against HNA-3a have been reported as one of the main causes of fatal TRALI in Germany in a few other countries ^25,26,27^. On the other hand, clinical consequences of alloantibodies against HNA-3b were found less severe.

In HNA-4 system, HNA-4a/a allele was found most common in India (90.18%) as seen in most global populations. Genotype frequency of HNA-4a was reported to range from 73%-100% ^16,23,28,18,21,15,14,17^ in various global population studies. Interestingly, in contrast to most global reports HNA-5b/b homozygous individuals represented 48.59% of the Indian population. HNA-5a/b, which was previously reported to be the most frequent in Zambians (52.5%)^21^ and Malaysians (49.5%)^15^ was found in about 38.09% of the population.

In conclusion, significant similarities albeit some differences in frequencies of HNA variants were observed in Indian population in comparison to other global populations. We sincerely hope these insights would guide cost-effective strategies for screening of HNAs. To the best of our knowledge, this is also the first and comprehensive study of HNA variants in the Indian population.

## Acknowledgements

MI acknowledges fellowship from ICMR-India. MD and BJ acknowledge research fellowships from CSIR-India. The funders had no role in the analysis of data, preparation of the manuscript or decision to publish.

## References

1. Browne T, Dearman RJ, Poles A. Human neutrophil antigens: Nature, clinical significance and detection. Int J Immunogenet. 2021;48(2):145–156.

2. Flesch BK, Reil A. Molecular Genetics of the Human Neutrophil Antigens. Transfusion Medicine and Hemotherapy. 2018;45(5):300–309. doi:10.1159/000491031

3. Lalezari P. Neutrophil-specific antigens, immunobiology, and implications in transfusion medicine and blood disorders. Transfusion. 2017;57(9):2066–2073.

4. Flesch BK, Curtis BR, de Haas M, Lucas G, Sachs UJ. Update on the nomenclature of human neutrophil antigens and alleles. Transfusion. 2016;56(6):1477–1479.

5. Lalezari P, Nussbaum M, Gelman S, Spaet TH. Neonatal neutropenia due to maternal isoimmunization. Blood. 1960;15:236–243.

6. Porcelijn L, de Haas M. Neonatal Alloimmune Neutropenia. Transfus Med Hemother. 2018;45(5):311–316.

7. Fung YL, Pitcher LA, Willett JE, et al. Alloimmune neonatal neutropenia linked to anti-HNA-4a. Transfus Med. 2003;13(1):49–52.

8. Mraz GA, Crighton GL, Christie DJ. Antibodies to human neutrophil antigen HNA-4b implicated in a case of neonatal alloimmune neutropenia. Transfusion. 2016;56(5):1161–1165.

9. Farruggia P, Dufour C. Diagnosis and management of primary autoimmune neutropenia in children: insights for clinicians. Ther Adv Hematol. 2015;6(1):15–24.

10. Akhtari M, Curtis B, Waller EK. Autoimmune neutropenia in adults. Autoimmun Rev. 2009;9(1):62–66.

11. Fujita M, Kawabata H, Oka T, et al. A Rare Case of Adult Autoimmune Neutropenia Successfully Treated with Prednisolone. Intern Med. 2017;56(11):1415–1419.

12. Bux J. Human neutrophil alloantigens. Vox Sang. 2008;94(4):277–285.

13. Esmaeili B, Bayat B, Alirezaee A, Delkhah M, Mehdizadeh MR, Pourpak Z. Human Neutrophil Antigen Genotype and Allele Frequencies in Iranian Blood Donors. J Immunol Res. 2022;2022:4387555.

14. Matsuhashi M, Tsuno NH, Kawabata M, et al. The frequencies of human neutrophil alloantigens among the Japanese population. Tissue Antigens. 2012;80(4):336–340.

15. Hajar CGN, Zulkafli Z, Md Riffin NS, et al. Human neutrophil antigen frequency data for Malays, Chinese and Indians. Transfus Apher Sci. 2020;59(2):102651.

16. Tam K, Tang I, Ho J, et al. A study of human neutrophil antigen genotype frequencies in Hong Kong. Transfus Med. 2018;28(4):310–318.

17. Changsri K, Tobunluepop P, Songthammawat D, Apornsuwan T, Kaset C, Nathalang O. Human neutrophil alloantigen genotype frequencies in Thai blood donors. Blood Transfus. 2014;12 Suppl 1:s286–s291.

18. Hauck B, Philipp A, Eckstein R, et al. Human neutrophil alloantigen genotype frequencies among blood donors with Turkish and German descent. Tissue Antigens. 2011;78(6):416–420.

19. Kissel K, Hofmann C, Gittinger FS, Daniels G, Bux J. HNA-1a, HNA-1b, and HNA-1c (NA1, NA2, SH) frequencies in African and American Blacks and in Chinese. Tissue Antigens. 2000;56(2):143–148.

20. Huvard MJ, Schmid P, Stroncek DF, Flegel WA. Frequencies of SLC44A2 alleles encoding human neutrophil antigen-3 variants in the African American population. Transfusion. 2012;52(5):1106–1111.

21. Nielsen KR, Koelbaek MD, Varming K, Baech J, Steffensen R. Frequencies of HNA-1, HNA-3, HNA-4, and HNA-5 in the Danish and Zambian populations determined using a novel TaqMan real time polymerase chain reaction method. Tissue Antigens. 2012;80(3):249–253.

22. Karczewski KJ, Francioli LC, Tiao G, et al. The mutational constraint spectrum quantified from variation in 141,456 humans. Nature. 2020;581(7809):434–443.

23. Xia W, Bayat B, Sachs U, et al. The frequencies of human neutrophil alloantigens in the Chinese Han population of Guangzhou. Transfusion. 2011;51(6):1271–1277.

24. Reil A, Wesche J, Greinacher A, Bux J. Geno- and phenotyping and immunogenicity of HNA-3. Transfusion. 2011;51(1):18–24.

25. Keller-Stanislawski B, Reil A, Günay S, Funk MB. Frequency and severity of transfusion-related acute lung injury--German haemovigilance data (2006-2007). Vox Sang. 2010;98(1):70–77.

26. Reil A, Keller-Stanislawski B, Günay S, Bux J. Specificities of leucocyte alloantibodies in transfusion-related acute lung injury and results of leucocyte antibody screening of blood donors. Vox Sang. 2008;95(4):313–317.

27. Davoren A, Curtis BR, Shulman IA, et al. TRALI due to granulocyte-agglutinating human neutrophil antigen-3a (5b) alloantibodies in donor plasma: a report of 2 fatalities. Transfusion. 2003;43(5):641–645.

28. Cardoso SP, Chong W, Lucas G, Green A, Navarrete C. Determination of human neutrophil antigen-1, -3, -4 and -5 allele frequencies in English Caucasoid blood donors using a multiplex fluorescent DNA-based assay. Vox Sang. 2013;105(1):65–72.

